# Anti-TGFβ peptide P144 augments CAR-T cell efficacy through neutrophil recruitment and TME remodeling in malignant mesothelioma

**DOI:** 10.64898/2026.10.28.762702

**Authors:** Reyisa Bughda, Elizabeth Hawkins, Gary Box, Florencia Secchiari, Clara Leboreiro Babé, Ray S Tan, Vivian Dimou, Reena R D’Souza, Astero Klampatsa

## Abstract

Malignant mesothelioma is a rare, aggressive and highly desmoplastic cancer characterised by a profoundly immunosuppressive tumour microenvironment (TME). Chimeric antigen receptor (CAR)-T cell therapy has transformed the treatment of hematological malignancies, but its activity against solid tumours such as mesothelioma remains limited, in part through transforming growth factor-beta (TGFβ)-mediated immunosuppression. To address this barrier, we engineered murine CAR-T cells that simultaneously target tumour antigen and deliver TGFβ blockade to the tumour. We first compared two anti-TGFβ peptides, the beta glycan-derived P144 and the phage-display-derived P17, in vitro; both attenuated TGFβ-induced epithelial-to-mesenchymal transition and reporter activity, but P144 suppressed canonical Smad3 phosphorylation more effectively and was selected for downstream work. Murine CAR constructs targeting either mesothelin (M11) or fibroblast activation protein (FAP) were modified to secrete P144, and the resulting products were assessed in co-culture assays and in the syngeneic AE17m model of mesothelioma. P144 secretion did not alter transduction efficiency, expansion or antigen-specific cytotoxicity, but markedly increased antigen-driven IFNγ production, most notably at low effector-to-target ratios. In vivo, both M11-P144 and FAP-P144 CAR-T cells-controlled tumour growth better than their parental, peptide-free counterparts, although complete tumour resolution was not achieved. Flow cytometric profiling of the tumour immune infiltrate showed that P144 delivery increased intratumoral neutrophil frequency approximately three-fold, and neutrophils recovered from these tumours were substantially more cytotoxic ex vivo than those from control tumours. Antibody-mediated neutrophil depletion attenuated the additional benefit conferred by P144 while leaving parental CAR-T activity largely intact, indicating that neutrophils contribute functionally to the effect. Local, CAR-T-delivered TGFβ blockade therefore remodels the mesothelioma TME and engages a tumoricidal neutrophil response, offering a modular strategy for enhancing adoptive cell therapy in desmoplastic solid tumours.

## Introduction

Malignant mesothelioma is a rare, highly aggressive and desmoplastic malignancy of mesothelial surfaces, predominantly linked to historical asbestos exposure (1–2). Despite multimodal clinical intervention — now including first-line immune checkpoint blockade for unresectable disease — the prognosis remains dismal, in large part because of a dense and profoundly immunosuppressive tumour microenvironment (TME) (1–5).

Transforming growth factor-beta (TGFβ) is a pleiotropic cytokine with a dual role in cancer biology: it restrains tumour growth at early stages, but during later disease progression, it promotes invasion, metastasis and immune evasion (6–7). In mesothelioma, TGFβ is frequently upregulated within the TME, where it drives proliferation of cancer-associated fibroblasts and pushes tumour cells towards a more invasive phenotype through epithelial-to-mesenchymal transition (EMT) (1–2, 8–9). Dysregulated TGFβ signaling correlates with shorter patient survival and with resistance to standard therapies, marking it as a driver of poor clinical outcome (10). Systemic pharmacological blockade of TGFβ has proven difficult, both because of the complexity of the signaling network and because TGFβ is required for systemic homeostasis and peripheral immune tolerance. On-target toxicity has thus limited the clinical development of several pathway inhibitors (11–13). Attention has therefore shifted towards localized, immune-based combination strategies, and approaches under investigation include bifunctional checkpoint–TGFβ traps, selective blockade of latent TGFβ activation and cell-mediated delivery of TGFβ antagonists (13).

Chimeric antigen receptor (CAR)-T cell therapy has transformed the management of hematological malignancies but remains of limited benefit in solid tumours (14–16), and in mesothelioma its activity is constrained both by physical desmoplastic barriers and by the immunosuppressive milieu that local TGFβ production sustains (1–2). Co-targeting strategies that pair a tumour-associated antigen with TGFβ blockade confined to the tumour are consequently attractive. Two targets are of particular interest in mesothelioma: mesothelin, a cell-surface glycoprotein overexpressed on mesothelial malignancies (17–19), and fibroblast activation protein (FAP), which is enriched on the surrounding tumour stroma (20–21).

Here, we examined whether the secretory peptide TGFβ inhibitors P17 and P144 could be used as modular components to relieve TGFβ-mediated suppression directly within the mesothelioma TME. P144 is derived from the ligand-binding domain of the TGFβ type III receptor (beta glycan), whereas P17 was identified by phage-display screening; both have been shown to block TGFβ signaling, limit tumour-induced immune evasion and improve T-cell responsiveness (22–24). We first compared the two candidates *in vitro* and identified P144 as the more potent inhibitor of canonical Smad3 signaling. We then engineered murine CAR constructs targeting either mesothelin or FAP to constitutively secrete P144. Using the syngeneic AE17 mesothelioma model, we show that these dual-function constructs outperform conventional CAR-T cells in controlling tumour progression, and that local P144 delivery remodels the myeloid compartment by mobilizing a population of cytotoxic, tumoricidal neutrophils that are functionally required for the additional therapeutic benefit.

## Materials and Methods

### Cell lines

Phoenix packaging cells, the human mesothelioma line EM, the human lung cancer A459 line, the murine mesothelioma line AE17, and the 3T3 murine fibroblast cell line were kindly provided by Prof. Steven Albelda (University of Pennsylvania, USA) and maintained in RPMI-1640 supplemented with 10% FBS, 1% penicillin–streptomycin and 1% L-glutamine (all Gibco). EM and AE17 cells had previously been lentivirally transduced to stably express human mesothelin, generating EMMESO and AE17m respectively (25). 3T3 had been transduced to express human FAP, generating the 3T3-FAP cell line (26). The HEK-Blue™ TGFβ reporter line (InvivoGen) was cultured in DMEM supplemented with 10% FCS, 1% penicillin–streptomycin, 1% L-glutamine and 1% HEPES (all Gibco), together with blasticidin (10 μg/ml), hygromycin B gold (100 μg/ml), Zeocin™ (100 μg/ml) and Normocin™ (50 μg/ml) (all InvivoGen). All lines were maintained at 37 °C in 5% CO₂.

### Epithelial-to-mesenchymal transition (EMT) assay

EMMESO or A549 cells (3 × 10⁵) were seeded into six-well plates and allowed to adhere for 4 h at 37 °C in 5% CO₂. Peptides P17 (KRIWFIPRSSWYERA) and P144 (TSLDASIIWAMMQN) were synthesized and delivered by Genscript. The peptides were reconstituted at 1 mg/mL in water and sonicated for 15 minutes at 20 % amplitude. Cells received two doses of recombinant TGFβ (5ng/ml), P17 (100 μg/ml) or P144 (100 μg/ml), alone or in combination, over two days. Cells were harvested in PBS, centrifuged (1,500 rpm, 5 min), resuspended in 100 μl PBS and stained with Live/Dead Blue viability dye (Thermo Fisher Scientific, 1:600) for 30 min at 4 °C. After washing, cells were stained with anti-E-cadherin-APC (BioLegend, 2 μl/sample) and anti-EpCAM-PE Dazzle 594 (BioLegend, 2 μl/sample) for 30 min at 4 °C. Samples were acquired on a BD Symphony A5 cytometer and analysed in FlowJo v10.10.0 and GraphPad Prism v11.0.2.

### TGFβ reporter assay

HEK-Blue™ TGFβ reporter cells were seeded at 5 × 10⁴ cells per well in flat-bottomed 96-well plates and treated with recombinant TGFβ (5ng/ml), P17 (100 μg/ml) or P144 (100 μg/ml), alone or in combination, for 24 h at 37 °C in 5% CO₂. Plates were centrifuged (1,300 rpm, 5 min) and 20 μl of supernatant per well transferred to a fresh plate containing 180 μl QUANTI-Blue™ reagent (InvivoGen). Absorbance was read at 635 nm at the indicated timepoints and analysed in GraphPad Prism.

### TGFβ ELISA

TGF-β1 secreted by human and murine cell lines were determined by measuring its concentration in the culture supernatant. Human cell lines used were EMMESO, HT1080, and REN. Murine cell lines used were AE17 and AB1. The cells were cultured as described before until they reached confluency. They were then washed 3x with PBS and switched to serum free RPMI media for up to 48 hours. Supernatants were collected one-week post-culture, and the TGF-β1 concentration was determined using the Human/Mouse/Rat/Porcine/Canine TGF-beta 1 ELISA Kit (R&D systems) and following the manufacturer’s protocol. Optical density was read at 450 nm and analysed in GraphPad Prism v11.0.2.

### Western blotting

EMMESO cells collected from the EMT assay were pelleted and lysed using 1 x RIPA buffer (Millipore) in the presence of Halt™ Protease and Phosphatase Inhibitor Cocktail (Thermo Fisher), incubated on ice for 1 hour. The mixture was then centrifuged at 10,000 x g for 10 minutes to pellet cell debris. The supernatant was transferred to a fresh tube, total protein extracted in the supernatant was quantified using Pierce™ BCA Protein Assay Kit. The supernatant was diluted in PBS and plated in 96 well microplate along with Diluted Albumin Standards in appropriate wells. The samples were incubated with 200 μL Working Reagent for 30 minutes at 37 °C. Absorbance was measured at 562 nm on a plate reader. Concentration of total protein in the supernatant was calculated from extrapolation of standard curve. 5 μg of protein extracted from each treatment condition was denatured using 2-Mercaptoethanol (Sigma-Adrich) diluted in LaemmLi sample Buffer (BioRad) and heated at 95 °C_for 5 minutes, the samples and a protein standard were loaded onto 4-15 % MINI-PROTEAN® Protein Gels for SDS-PAGE to separate proteins based on their molecular weight. Electrophoresis was performed at 200 V for 30 minutes, the separated proteins on the gel were transferred to polyvinylidene difluoride (PVDF) membrane, using Trans-Blot Turbo Transfer System (Bio-Rad). Following the transfer, the membrane was placed in 5 % PBS to block overnight at 4 °C. The membrane was washed three times using 0.05 % Tween 20 in PBS, incubating the membrane in wash buffer for 10 minutes each time. The membrane was incubated in primary antibodies (recombinant anti-SMAD3, 1:2,000; anti-β actin, 1:2,000; both BioRad) for 1 hour at RT and washed again for three times before incubating with secondary antibodies for 1 hour. The membrane was visualized in G: BOX (Syngene) imaging system. For β-actin staining, the primary and secondary antibodies were stripped off from the membrane by incubating in Restore™ Western Blot Stripping Buffer (ThermoFisher) for 15 minutes at 37°C. The membrane was washed three times and incubated with anti-β actin antibody for 1 hour at RT. Secondary antibody was applied for 1 hour following another round of washing before imaging. Intensities of the bands were determined using ImageJ software, the values were normalized by diving the band intensity by corresponding loading control (β-actin) intensity.

### Generation of CAR constructs and primary murine T cells

The anti-mesothelin M11 CAR construct (25) and anti-fibroblast activation protein (FAP) CAR construct (26), kindly provided by Steven M. Albelda (University of Pennsylvania), were modified to constitutively secrete the TGFβ inhibitory peptide P144. The published P144 sequence (TSLDASIIWAMMQN), derived from residues 730-743 of the TGFβ type III receptor (beta glycan) (27), was cloned downstream of an IL-2 secretion signal and upstream of GFP. To enable co-expression, P144 and GFP were separated by a self-cleaving T2A peptide. In both constructs, the original IRES-GFP cassette was replaced with an IRES– IL2 signal peptide–P144–2A–GFP cassette, generating the M11-P144 and FAP-P144 CAR vectors shown in Figures 3A and 5A. Infective particles were generated from the supernatants of Phoenix cells transfected with the retroviral vector plasmid and helper plasmids using Lipofectamine 2000 (Invitrogen) (28). Primary murine splenic CD3⁺ T cells were isolated from naïve C57BL/6 mice, activated and retrovirally transduced following a previously optimised protocol (25).

### In vitro cytotoxicity and IFNγ assays

Triplicate wells containing 5,000 luciferase-expressing AE17m cells (mesothelin-targeted experiments) or 3T3-FAP cells (FAP-targeted experiments) were co-cultured with the indicated CAR-T cells at a range of effector-to-target (E:T) ratios. Cytotoxicity was measured following overnight co-culture using the luciferase assay system (Promega) as previously described (25). IFNγ in co-culture supernatants was quantified with a mouse IFNγ ELISA kit (BioLegend) (25).

### Syngeneic tumour models and in vivo therapy

Female C57BL/6 mice (6–8 weeks, 16–18 g) were purchased from Charles River Laboratories (UK). AE17m cells (2 × 10⁶) were injected subcutaneously into the right flank. Treatment began when tumours reached approximately 50–80 mm³, around five days after inoculation. Tumour volumes and body weights were recorded every three to four days until the regulatory endpoint.

#### Study 1 — CAR-T cells combined with free peptide (*Figure 2*)

Mice were allocated to one of seven groups: untreated; non-transduced (NTD) T cells; P144 alone; M11 CAR-T cells; or M11 CAR-T cells plus P144. CAR-T cells (1 × 10⁷ transduced cells) were given as a single intravenous tail-vein injection on day 5. Peptides were administered intratumorally at 100 μg in 50μl PBS on days 5, 7, 9, 12 and 14, and tumours were harvested on day 15.

#### Study 2 — P144-secreting CAR-T cells (*Figure 3*)

Mice were allocated to untreated, NTD, M11 or M11-P144 groups and received a single intravenous injection of 1 × 10⁷ transduced cells on day 5. Tumours were harvested on day 18.

#### Study 3 — Neutrophil depletion (*Figure 4F*)

Tumour-bearing mice received intraperitoneal anti-Ly6G antibody (clone 1A8, BioXCell; 100 μg per mouse) one day before CAR-T infusion and twice weekly thereafter for up to two weeks. [AUTHOR ACTION: state the isotype control used in the wild-type arm, and report the depletion efficiency achieved, measured by flow cytometry on blood or tumour.]

#### Study 4 — FAP-targeted CAR-T cells (*Figure 5E*)

Mice were allocated to untreated, NTD, FAP or FAP-P144 groups and treated as in Study 2, with tumours monitored to day 20. [AUTHOR ACTION: confirm cell dose and route if these differed from Study 2.]

### Flow cytometry

At the experimental endpoint, tumours and tumour-draining lymph nodes (TDLNs) were harvested and processed into single-cell suspensions. Tumours were minced and digested in a 1:1 mixture of L-15 medium (Gibco) and digestion buffer as previously described (25). Digested tumours and intact TDLNs were passed through 70-μm strainers, centrifuged at 1,500 rpm for 5 min and subjected to red blood cell lysis (Gibco) where required. Suspensions were washed, filtered and resuspended in PBS. Samples were divided between T-cell and myeloid panels. Cells were incubated with Live/Dead UV450/50 viability dye (Invitrogen, 1:600) and mouse Fc block (BD Biosciences, 1:200) for 15 min at 4 °C, washed, and stained with surface antibody master mixes (1:200) for 40 min at 4 °C. For intracellular markers, cells were fixed and permeabilized using the eBioscience kit according to the manufacturer’s instructions and stained with intracellular antibody master mixes (1:200) for 1 hr at 4 °C. After final washes, cells were fixed in 4% paraformaldehyde, resuspended in PBS and stored at 4 °C until acquisition. Antibodies used were: CD45 (30-F11, Biolegend), CD3 (17A2, Biolegend), CD8 (53-6.7, BD Biosciences), CD4 (GKI.5, Biolegend), CD11b (M1/70, Biolegend), Ly6G (1A8, Biolegend), IFNγ (XMG1.2, BD Biosciences), Ki67 (16A8, Biolegend), CD25 (PC61, Biolegend), Ly6C (HK1.4, Biolegend), F4/80 (F45-2342, BD Biosciences), CD206 (C068C2, Biolegend), NK1.1 (PK136, Biolegend).

### Neutrophil isolation and cytotoxicity assay

Tumours were dissociated into single-cell suspensions and neutrophils isolated using the EasySep FITC Positive Selection Kit II (StemCell Technologies). Suspensions were incubated with mouse Fc block (BD Biosciences), stained with FITC-conjugated anti-Ly6G antibody (BioLegend) for 15 min at room temperature, then incubated sequentially with the selection cocktail and RapidSpheres (StemCell Technologies) before positive selection on an EasySep magnet. AE17m cells were seeded at 5,000 cells per well in 96-well plates and allowed to adhere for 4 h. Freshly isolated tumour neutrophils were added at the indicated E:T ratios and co-cultured for 24 h. Wells were washed with PBS, replenished with medium and treated with an equal volume of CellTiter-Glo reagent (Promega). Plates were shaken for 5 min, incubated for 10 min at room temperature, and luminescence read on a plate reader. Data were normalized to untreated control wells and analysed in GraphPad Prism.

### Statistical analysis

Analyses were performed in GraphPad Prism (v7-11). Comparisons were made using the non-parametric Mann–Whitney test, one-way ANOVA with Dunnett’s multiple comparisons test, or two-way ANOVA, as appropriate to the design and as indicated in each figure legend. p < 0.05 was considered significant. Variability is presented as the standard error of the mean (SEM) throughout, and n refers to the number of independent biological replicates or individual animals as stated in each legend.

## Results

### P17 and P144 partially attenuate TGFβ-induced epithelial-to-mesenchymal transition

To determine whether P17 and P144 retained biological activity, we first tested their capacity to counteract TGFβ-induced EMT. EMT was induced with recombinant TGFβ and read out as surface expression of the epithelial markers E-cadherin and EpCAM in A549 lung adenocarcinoma and EMMESO mesothelioma cells (Figure 1A). In A549 cells, TGFβ markedly reduced expression of both markers. E-cadherin was significantly reduced relative to untreated controls (p = 0.0007), and co-treatment with either peptide partially restored it, yielding significantly higher expression than TGFβ alone (p = 0.0011 for P17 and p = 0.0019 for P144). TGFβ similarly reduced EpCAM expression from 84.5% to 22.7%, while co-treatment with P17 or P144 restored expression to 75.0% and 69.7% respectively. Neither peptide alone altered marker expression relative to untreated controls. EMMESO cells behaved comparably (Figure 1A). TGFβ reduced E-cadherin from 87.9% to 28.8% and EpCAM from 70.8% to 23.4%. Co-treatment preserved expression substantially, restoring E-cadherin to 84.2% with P17 and 76.5% with P144, and EpCAM to 61.9% and 59.4% respectively. As in A549 cells, neither peptide affected marker expression in the absence of TGFβ. Both peptides therefore attenuate TGFβ-driven EMT in epithelial and mesothelial tumour models.

**Figure 1.**
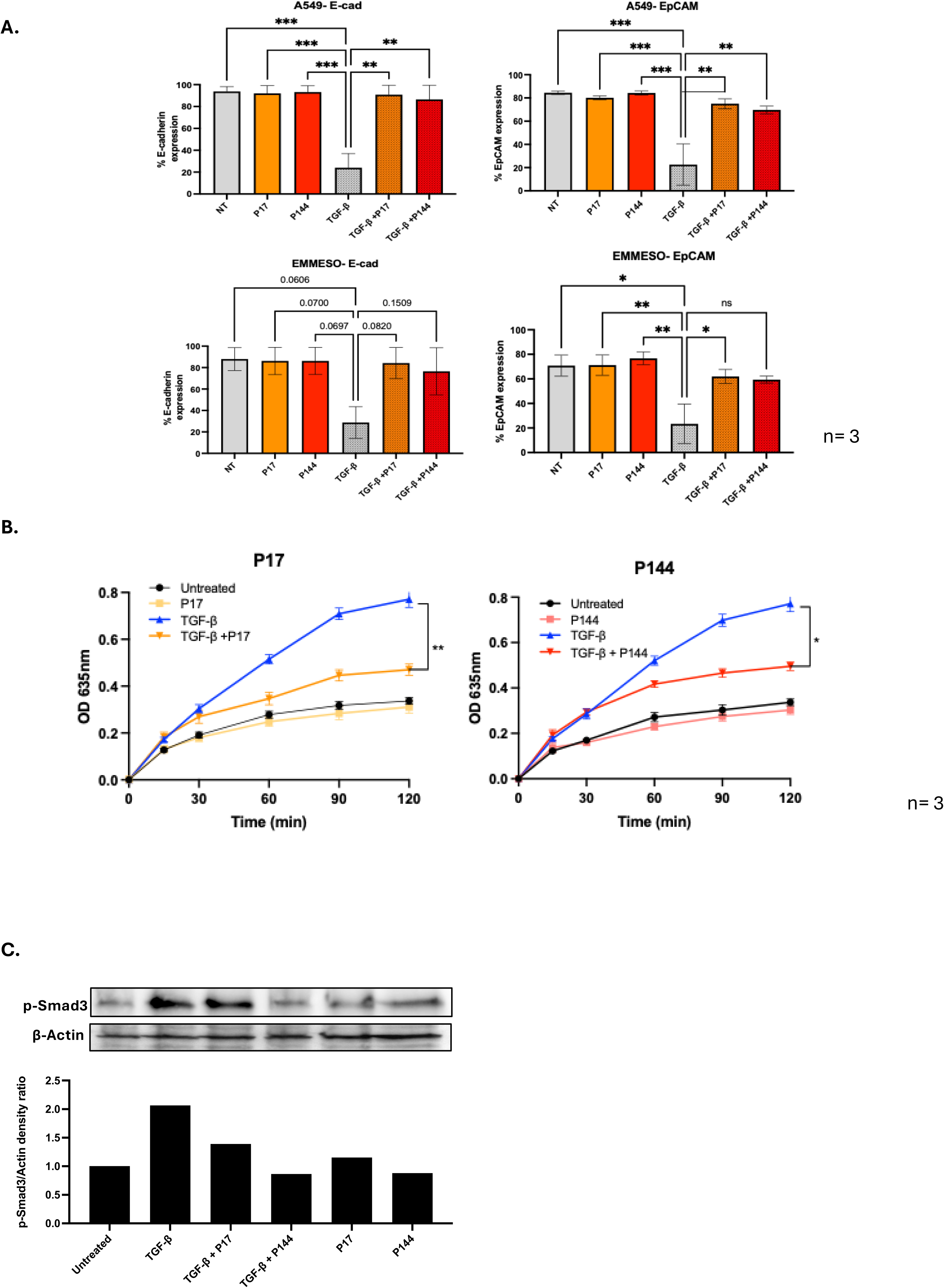
P17 and P144 attenuate TGF-β signalling and EMT in vitro. **(A)** A549 lung adenocarcinoma and EMMESO mesothelioma cells were cultured in the presence or absence of TGF-β and treated with either P17 or P144. Expression of the epithelial markers E-cadherin and EpCAM was assessed to evaluate epithelial-to-mesenchymal transition (EMT). Both peptides partially preserved epithelial marker expression following TGF-β stimulation. **(B)** HEKBlue TGF-β reporter cells were stimulated with TGF-β in the presence or absence of P17 or P144. Reporter activity was measured over time by optical density at 635 nm. Both peptides reduced TGF-β-induced reporter activation while having minimal effects on basal signalling. **(C)** Canonical TGF-β signalling was assessed by Western blot analysis of phosphorylated Smad3 (p-Smad3). β-actin served as a loading control. Representative blots and densitometric quantification are shown. P144 produced greater inhibition of Smad3 phosphorylation than P17 and was selected for incorporation into subsequent CAR-T cell constructs. Data are presented as indicated in the individual panels. Statistical analyses were performed as described in the Materials and Methods. Statistical significance is indicated as follows: *P* < 0.05, **P** < 0.01, ***P*** < 0.001 and **P** < 0.0001; ns, not significant.

### Both peptides inhibit TGFβ signalling in a reporter assay

We next asked whether these effects reflected inhibition of canonical TGFβ signalling. HEK-Blue TGFβ reporter cells were stimulated with TGFβ in the presence or absence of each peptide and reporter activity followed over 120 min (Figure 1B). TGFβ produced a progressive rise in reporter activity, reaching an OD₆₃₅ of 0.771 at 120 min. Neither peptide alone altered basal activity. Both significantly attenuated TGFβ-induced signalling: P17 reduced reporter activity to 0.470 at 120 min, a 39% reduction relative to TGFβ alone (p<0.01), and P144 to 0.496, a 36% reduction (p<0.01). These data confirm that both peptides suppress TGFβ pathway activity directly and suggest that the preservation of epithelial marker expression above is mediated through TGFβ pathway inhibition.

### P144 suppresses canonical Smad3 activation more effectively than P17

To compare the potency of the two peptides, we examined phosphorylation of Smad3, the principal downstream mediator of canonical TGFβ signalling (Figure 1C). TGFβ stimulation increased the p-Smad3/β-actin ratio to 1.79-fold relative to untreated controls. Co-treatment with P17 reduced this only partially, to 1.43, whereas P144 suppressed it to 0.88, close to baseline. Relative to TGFβ-treated controls, this corresponds to reductions of approximately 20% for P17 and 51% for P144. Neither peptide substantially altered basal Smad3 phosphorylation in the absence of TGFβ (P17, 0.96; P144, 0.71). Taken with the reporter data, these findings show that both peptides inhibit TGFβ signalling but that P144 blocks canonical Smad3 activation more effectively.

### P144 enhances the antitumour activity of M11 CAR-T cells in vivo

Having identified P144 as the more potent inhibitor of canonical signalling in vitro, we asked whether its local inhibition of TGFβ could enhance the activity of mesothelin-targeted CAR-T cells in vivo. Mice bearing established AE17m tumours received M11 CAR-T cells intravenously on day 5post tumor inoculation, with or without intratumoral injection of P144 on days 5, 7, 9, 12 and 14. Control groups received peptide alone, non-transduced (NTD) T cells, or no treatment (Figure 2A). Baseline TGFβ release by AE17 cells, together with a panel of comparator lines, is shown in Supplementary Figure S1A and supports the use of this model for TGFβ-directed intervention. Untreated and NTD controls grew progressively, reaching mean volumes of ∼265mm³ and 223mm³ respectively by day 15 (Figure 2B). M11 CAR-T cells alone significantly restricted progression, to 92 mm³. Adding P144 produced near-complete growth arrest: volumes plateaued shortly after treatment and remained at 31.5 mm³ at day 15, a reduction of approximately 66% relative to M11 CAR-T cells alone and around 88% relative to untreated controls (statistics by ordinary one-way ANOVA) (Figure 2C). Peptide monotherapy was significantly less effective demonstrating modest, delayed growth, to ∼203mm³ on day 15. P144 therefore has limited activity on its own but substantially enhances tumour control when combined with CAR-T therapy. Since improved tumour control might reflect greater CAR-T accumulation, we quantified intratumoral CAR⁺ CD8 T cells at endpoint (Figure 2D). Tumours from mice treated with M11 CAR-T cells alone contained only 0.9% CAR⁺ cells, rising to 9.5% with P144, an approximately ten-fold, significant increase (Weltch’s t-test). Local TGFβ blockade thus promotes CAR-T accumulation and persistence within the tumour, which may contribute to the superior efficacy seen with P144. All treatments were well tolerated, with body weights remaining stable over the course of the study (Supplementary Figure S1B).

**Figure 2.**
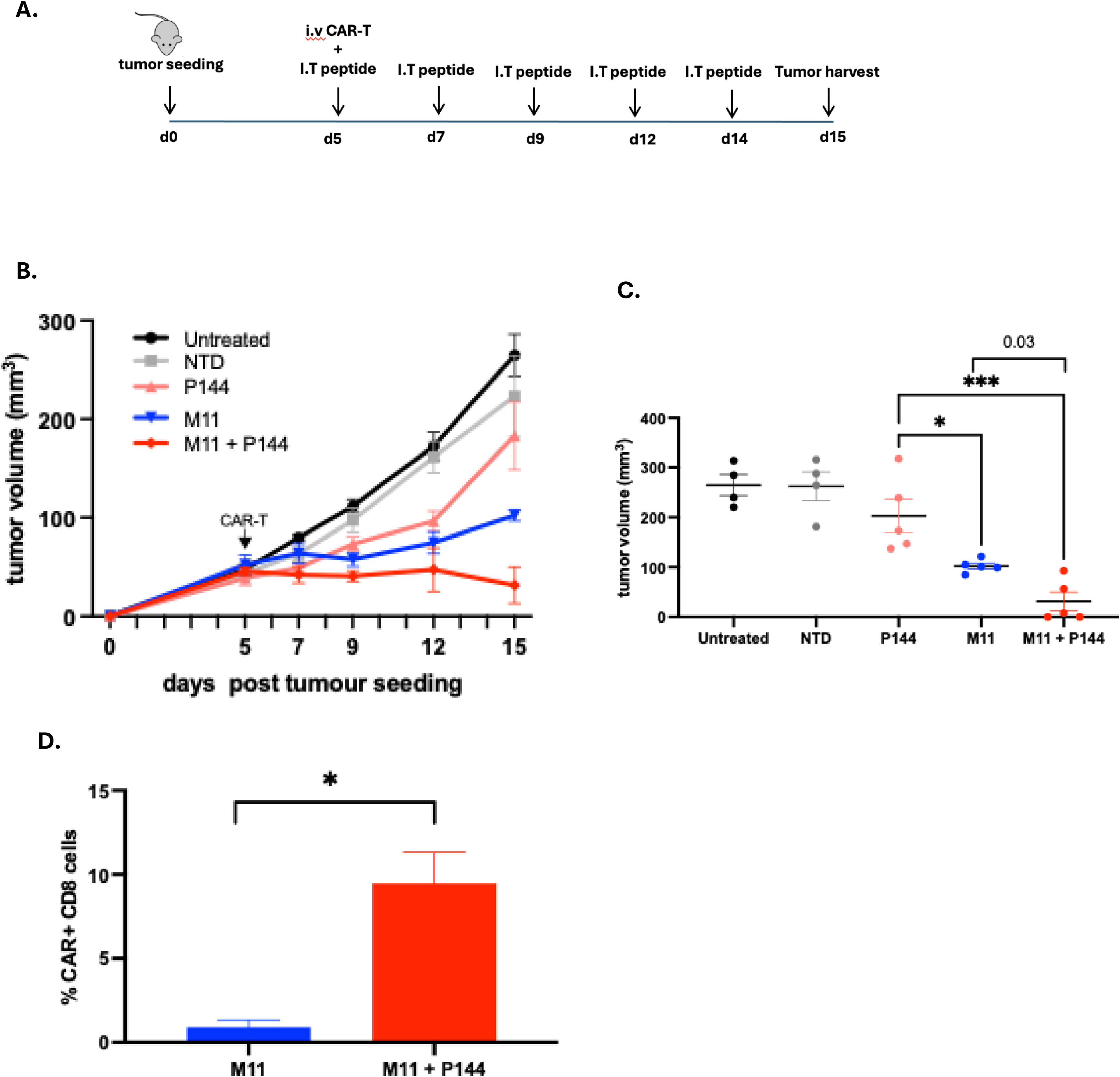
P144 enhances the antitumour efficacy of mesothelin-targeted CAR-T cells in vivo. **(A)** Experimental design. Tumour-bearing mice were treated with parental M11 CAR-T cells, M11 CAR-T cells combined with P17 or P144, non-transduced T cells, peptide alone, or left untreated. Tumour growth was monitored throughout the study and tumours were harvested at the experimental endpoint. **(B)** Tumour growth curves for the indicated treatment groups. Tumour volumes were measured serially using calipers and calculated as described in the Materials and Methods. Treatment with M11 CAR-T cells delayed tumour progression, while addition of P144 resulted in substantially improved tumour control. **(C)** Body weights recorded throughout the study. Body weight remained stable across all treatment groups, indicating that treatment was well tolerated. **(D)** Quantification of intratumoral CAR-positive T cells at study endpoint. Tumours were dissociated and analysed by flow cytometry. P144 treatment increased accumulation of CAR-positive T cells within the tumour microenvironment. Data are presented as indicated in the individual panels. Each symbol represents an individual animal where applicable. Statistical analyses were performed as described in the Materials and Methods.

### P144 secretion enhances CAR-T effector function and promotes enhanced tumor cell lysis in vivo

We next engineered the P144 cassette directly into the M11 CAR construct, placing it downstream of the IRES and upstream of a 2A-GFP element (Figure 3A), and compared the resulting M11-P144 product with parental M11 CAR-T cells. Both constructs transduced efficiently and comparably (Figure 3B, C). Mean CAR expression was ∼73% for M11 and 68% for M11-P144 across independent preparations, indicating that the secretion cassette does not impair transduction. Both products also expanded efficiently following activation, achieving approximately two-fold expansion over the culture period with no significant difference between groups (Figure 3D). Incorporating P144 therefore does not compromise manufacturing characteristics or ex vivo expansion. Antigen-specific cytotoxicity against mesothelin-expressing targets was likewise unaffected (Figure 3E). At an E:T ratio of 10:1, specific lysis reached 80% for M11 and 81% for M11-P144, and comparable killing was maintained at lower ratios. Cytokine production, by contrast, was markedly enhanced in the M11-P144 co-cultures (Figure 3F). At 10:1, IFNγ rose from 10,015 pg/ml to 11,431 pg/ml. The difference widened as target availability fell at 5:1, IFNγ increased from 5,436 to 7,750 pg/ml (p < 0.05), and at 1:1 from 1,336 to 6,008 pg/ml, an approximately 4.5-fold increase (p < 0.0001). Minimal cytokine was detected in non-transduced or untreated controls. Local TGFβ blockade therefore augments CAR-T effector function without altering antigen-specific killing capacity, a dissociation consistent with relief of a cytokine-production checkpoint rather than a change in cytolytic machinery. We then tested the M11-P144 construct in vivo (Figure 3G). Tumour growth was followed to day 18 (Figure 3H). Mice treated with parental M11 CAR-T cells reached a mean volume of 276 mm³ at endpoint, compared with 136mm³ for M11-P144, a reduction of approximately 51% (Figure 3J; p < 0.01). Both CAR-T groups were significantly smaller than untreated and NTD controls. Collectively, these results show that incorporating P144 into mesothelin-targeted CAR-T cells leaves transduction, expansion and cytotoxic function intact while significantly enhancing antigen-driven cytokine production and improving in vivo efficacy.

**Figure 3.**
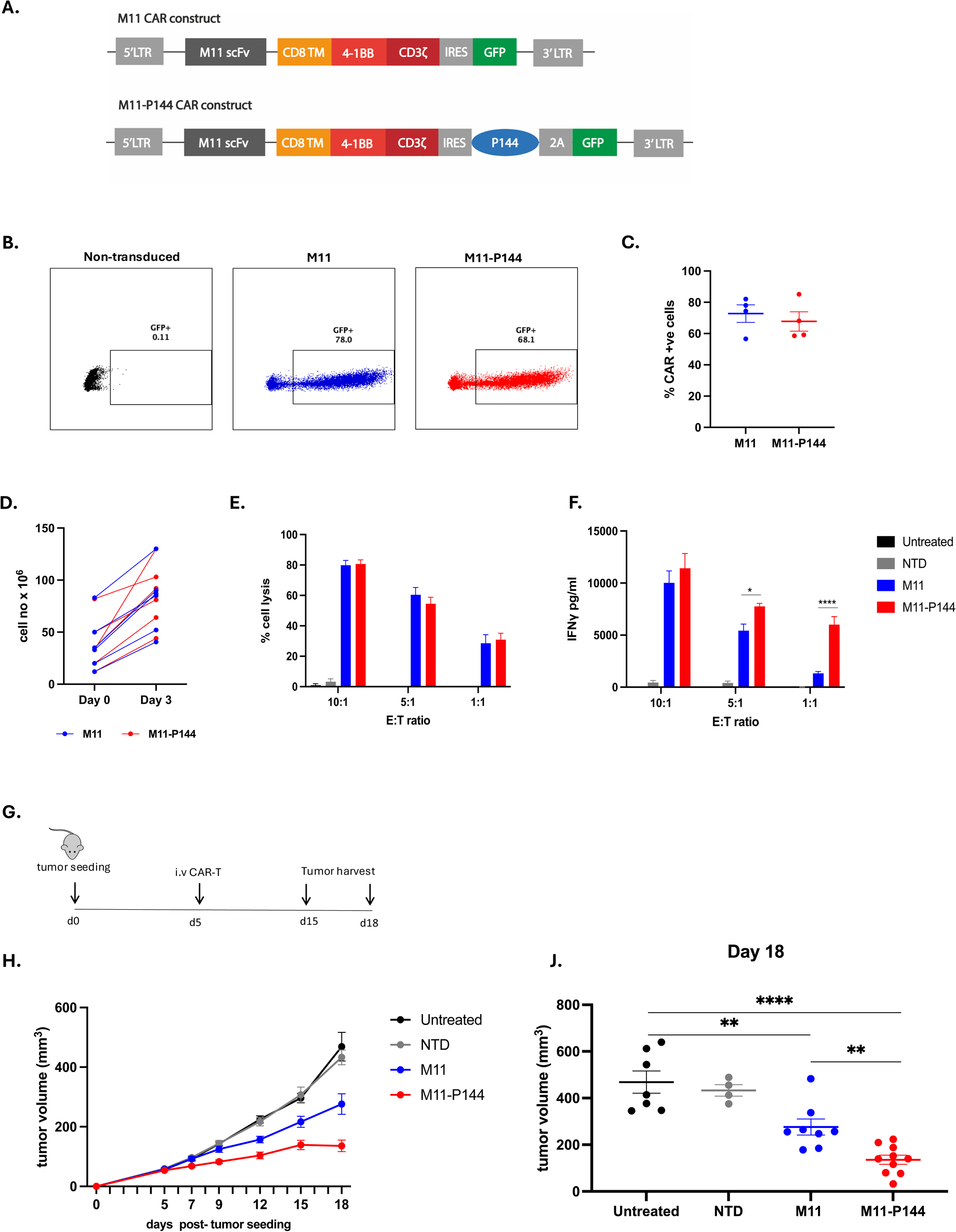
Engineering CAR-T cells to secrete P144 enhances effector function and antitumour activity. **(A)** CAR transduction efficiency in parental M11 and M11-P144 CAR-T cells. Representative flow cytometry plots and pooled donor data are shown. **(B)** Expansion of parental M11 and M11-P144 CAR-T cells following activation. Cell numbers were determined at the indicated time points and fold expansion was calculated relative to baseline. **(C)** Antigen-specific cytotoxicity of non-transduced, M11 and M11-P144 CAR-T cells against mesothelin-expressing target cells. Cytotoxicity was assessed at the indicated effector-to-target ratios. **(D)** IFN-γ secretion following co-culture with mesothelin-expressing target cells. Culture supernatants were collected and analysed by ELISA. M11-P144 CAR-T cells produced enhanced levels of IFN-γ compared with parental M11 CAR-T cells following antigen stimulation. **(E)** Independent in vivo validation study. Tumour-bearing mice were treated with parental M11 CAR-T cells or M11-P144 CAR-T cells and tumour growth was monitored over time. **(F)** Combined tumour growth data from three independent studies demonstrating the reproducibility of the enhanced antitumour activity associated with P144-secreting CAR-T cells. Data are presented as indicated in the individual panels. Statistical analyses were performed as described in the Materials and Methods. Statistical significance is indicated as follows: *P* < 0.05, **P** < 0.01, ***P*** < 0.001 and **P** < 0.0001; ns, not significant.

### P144 delivery increases intratumoral neutrophil frequency

To investigate the mechanisms underlying this enhanced activity, we characterised the tumour-infiltrating immune landscape by flow cytometry at endpoint. Analysis of the endogenous adaptive compartment showed a modest, non-significant increase in CD8⁺ T cell frequency in M11-P144-treated tumours relative to controls (Figure 4A), accompanied by numerically higher Ki67 and IFNγ expression among intratumoral CD8⁺ T cells (Figure 4B). A broader survey of T-cell and myeloid marker expression across treatment groups is provided in Supplementary Figures S1C and S1D. The myeloid compartment showed a clearer and more consistent change. Representative contour plots of CD11b⁺Ly6G⁺ gating are shown in Figure 4C, and quantification across groups in Figure 4D. Neutrophils, defined as CD45⁺CD11b⁺Ly6G⁺ cells, comprised approximately 2–3.5% of total live cells in untreated, non-transduced and M11-treated tumours, but rose to approximately 10% in M11-P144-treated tumours, a significant enrichment relative to all three comparator groups (p < 0.05). A corresponding increase in Ly6G⁺ cells, alongside a reduction in M2-polarised macrophages across all treated groups, is evident in the broader myeloid panel (Supplementary Figure S1D).

**Figure 4.**
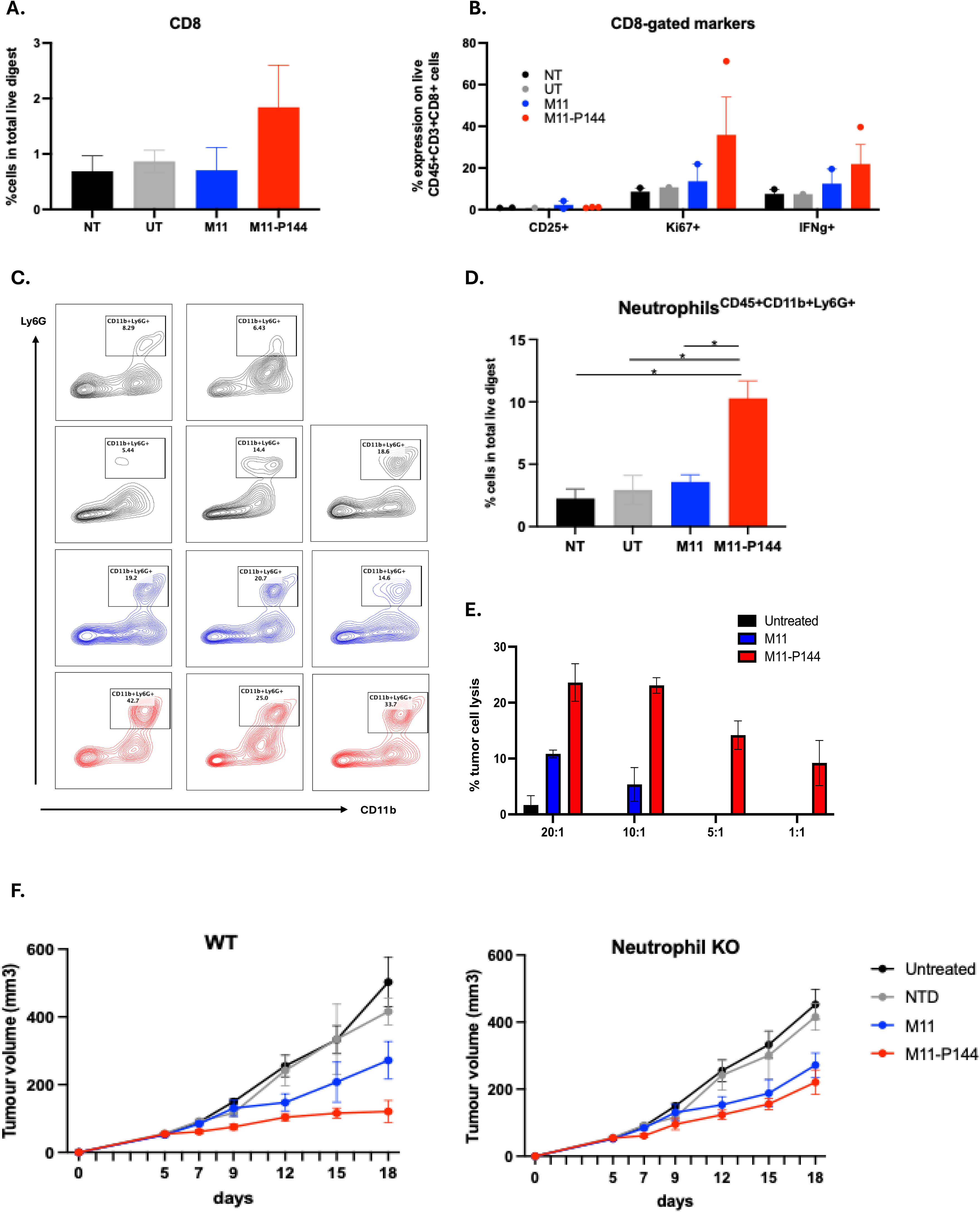
P144-mediated TGF-β blockade remodels the tumour microenvironment and promotes neutrophil-dependent antitumour activity. **(A)** Representative flow cytometric analysis of tumour-infiltrating neutrophils. Neutrophils were identified as live CD45⁺CD11b⁺Ly6G⁺ cells following exclusion of debris, doublets and dead cells. Representative gating strategy and quantification are shown. **(B)** Cytotoxic activity of tumour-derived neutrophils. Neutrophils isolated from untreated, M11-treated or M11-P144-treated tumours were co-cultured with mesothelin-expressing target cells at the indicated effector-to-target ratios. Specific target cell lysis was quantified following co-culture. **(C)** Contribution of neutrophils to CAR-T cell efficacy in vivo. Wild-type (WT) or neutrophil-depleted (αLy6G-treated) tumour-bearing mice were treated with parental M11 or M11-P144 CAR-T cells. Tumour growth was monitored over time. Depletion of neutrophils reduced the enhanced antitumour activity observed with M11-P144 CAR-T cells, demonstrating a functional contribution of neutrophils to therapeutic efficacy. Data are presented as indicated in the individual panels. Statistical analyses were performed as described in the Materials and Methods. Statistical significance is indicated as follows: *P* < 0.05, **P** < 0.01, ***P*** < 0.001 and **P** < 0.0001; ns, not significant. WT, wild type; αLy6G, anti-Ly6G antibody; TME, tumour microenvironment.

### Tumour-infiltrating neutrophils from P144-treated tumours are more cytotoxic

To establish whether these infiltrating neutrophils possessed antitumour activity, tumour-derived neutrophils were isolated and tested for cytotoxicity ex vivo (Figure 4E). Neutrophils from untreated tumours showed minimal killing, and those from parental M11-treated tumours only modest activity. Neutrophils recovered from M11-P144-treated tumours, by contrast, killed tumour cells substantially more effectively. At a 20:1 E:T ratio they mediated 23.6% specific lysis, compared with 10.8% for M11-derived and 1.7% for untreated-derived neutrophils, and enhanced cytotoxicity was maintained at 10:1, 5:1 and 1:1, where control neutrophils showed little or no measurable activity. 3 replicate wells, n=1.

### Neutrophils contribute functionally to the efficacy of P144-secreting CAR-T cells

To test whether neutrophils contribute directly rather than incidentally, tumour-bearing mice were treated with parental M11 or M11-P144 CAR-T cells with or without antibody-mediated neutrophil depletion (Figure 4F). In wild-type mice, M11-P144 CAR-T cells again controlled tumour growth better than parental M11 cells, reaching approximately 120 mm³ versus 270 mm³ at day 18. Following neutrophil depletion, this advantage was substantially attenuated, with M11-P144-treated tumours reaching approximately 220 mm³ while parental M11 activity was essentially unchanged at approximately 270 mm³. Neutrophils are therefore largely dispensable for baseline CAR-T-mediated tumour control but become an important effector arm specifically when TGFβ signalling is locally blocked. We note that the depletion attenuated rather than abolished the P144 benefit, indicating that neutrophils account for a substantial part, but not all, of the additional efficacy.

### FAP-P144 CAR-T cells are efficiently manufactured and retain antigen-specific function

Having established that mesothelin-targeted M11-P144 CAR-T cells involve neutrophils to enhance efficacy, we asked whether the strategy generalises to a stroma-directed construct. We generated FAP-targeted CAR-T cells carrying the same P144 secretion cassette (Figure 5A) and compared them with parental FAP CAR-T cells. Both constructs transduced efficiently, with GFP expression of 72.5% for FAP and 88.4% for FAP-P144 (Figure 5B), and both expanded substantially between days 2 and 5 of culture relative to non-transduced controls, with comparable proliferation between constructs (Figure 5C). When reacted with 3T3-FAP target cells, both products showed significant antigen-specific cytotoxicity and IFNγ secretion relative to non-transduced controls across all E:T ratios tested (Figure 5D). As with the mesothelin-targeted construct, FAP-P144 CAR-T cells produced significantly higher IFNγ levels than parental FAP CAR-T cells (statistics by one-way ANOVA)

**Figure 5.**
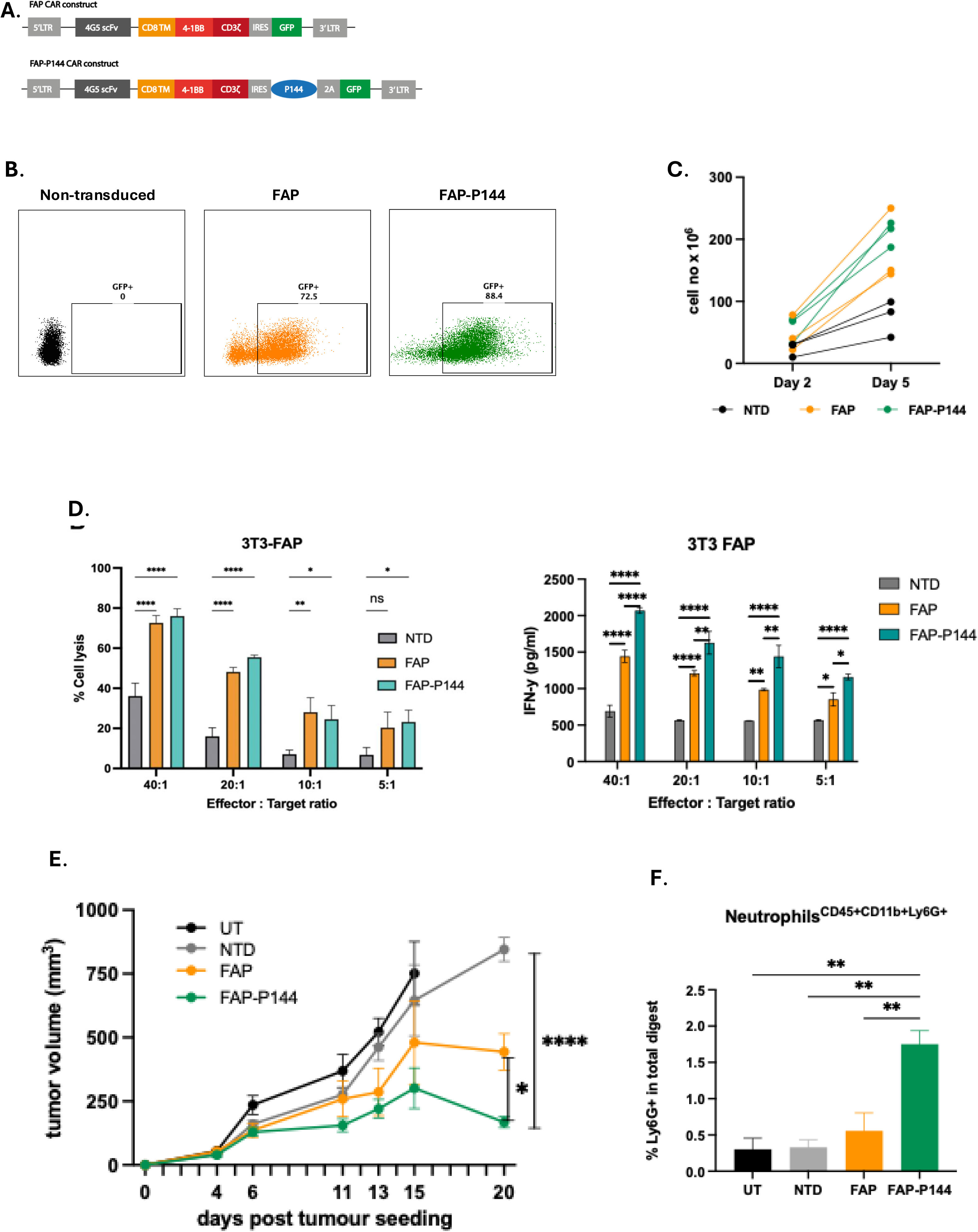
P144 secretion enhances the activity of FAP-targeted CAR-T cells. **(A)** Schematic representation of the parental FAP-targeted CAR construct and the P144-secreting FAP CAR construct. **(B)** Representative flow cytometry plots showing transgene expression in non-transduced, FAP and FAP-P144 CAR-T cells. **(C)** Quantification of transduction efficiency in the indicated CAR-T cell products. **(D)** Antigen-specific cytotoxicity against FAP-expressing target cells assessed at the indicated effector-to-target ratios. **(E–F)** Functional characterisation of parental FAP and FAP-P144 CAR-T cells following target recognition. Incorporation of the P144 secretion cassette preserved CAR-dependent function and enhanced responses against FAP-expressing targets. Data are presented as indicated in the individual panels. Statistical analyses were performed as described in the Materials and Methods. Statistical significance is indicated as follows: *P* < 0.05, **P** < 0.01, ***P*** < 0.001 and **P** < 0.0001; ns, not significant.

### FAP-P144 CAR-T cells outperform parental FAP CAR-T cells and recruit neutrophils in vivo

We next evaluated the activity of FAP-P144 CAR-T cells in vivo over a 20-day study (Figure 5E). FAP-P144-treated tumours grew significantly more slowly than untreated and non-transduced controls (p < 0.0001) and also outperformed parental FAP CAR-T cells (p < 0.05). On experimental endpoint day 20 post tumour seeding, mice treated with FAP-P144 reached a mean volume of 168 mm^3^, compared with mice treated with FAP CAR-T cells that had a mean volume of 444 mm^3^ and non-transduced T cells with mean volume of 845 mm^3^ respectively. Consistent with the mesothelin-targeted model, flow cytometric analysis showed a significant increase in CD45⁺CD11b⁺Ly6G⁺ neutrophil infiltration in FAP-P144-treated tumours relative to untreated, non-transduced and parental FAP CAR-T-treated tumours (Figure 5F; p < 0.01). P144-mediated neutrophil recruitment and the accompanying gain in efficacy are therefore not restricted to mesothelin-targeted CAR-T cells, supporting the generalisability of this approach across CAR targeting modalities in mesothelioma.

## Discussion

This study demonstrates that CAR-T delivery of the anti-TGFβ peptide P144 enhances antitumour efficacy against malignant mesothelioma and identifies intratumoral neutrophils as an important contributor to that benefit. Both P144 and P17 peptides inhibited TGFβ signalling, with P144 more effectively suppressing Smad3 phosphorylation. CAR-T cells engineered to secrete P144 accumulated at higher frequencies within tumours and improved tumor control across mesothelin- and FAP-targeted platforms. Treatment also increased the abundance and cytotoxicity of tumor-infiltrating neutrophils, which were required for the full therapeutic effect. These findings support local, CAR-T delivered TGFβ blockade as a potentially modular strategy for overcoming the immunosuppressive and desmoplastic barriers that limit adoptive cell therapy in mesothelioma. Our in vitro comparisons confirmed that P17 and P144 attenuate TGFβ-induced EMT and canonical Smad-dependent signalling, consistent with their activity as direct TGFβ antagonists. P144 derives from the ligand-binding domain of the type III TGFβ receptor, beta glycan, whereas P17 was identified through phage-display screening (22,24,27). Originally developed as an antifibrotic peptide, P144 has demonstrated activity in multiple models of pathological fibrosis and tissue remodelling (27, 29–31). Its superior suppression of Smad3 phosphorylation in our system provided the rationale for further development in CAR-T cells. Mesothelioma is particularly relevant to this approach because its dense fibrotic stroma, elevated TGFβ activity and asbestos-associated EMT contribute to tumor progression and immune cell exclusion (1–2, 8). Our EMT and reporter assays demonstrated that TGFβ blockade can partially reverse these changes in mesothelioma and lung adenocarcinoma cells. Two in vivo observations support the therapeutic rationale. Combining M11 CAR-T cells with repeated intratumoral injections of free P144produced near complete growth arrest, whereas locally administered peptide alone achieved only modest delay (Figure 2). Engineering P144 secretion into the CAR construct subsequently improved tumor control by approximately 51% relative to the parental CAR-T cells in an independent study (Figure 3). Because free P144 was also delivered intratumorally, the combination benefit cannot be explained simply by improved peptide penetration; rather, it appears to depend on the simultaneous presence of antigen-specific effector cells. These findings add evidence that TGFβ blockade can enhance CAR-T cell activity. Dominant-negative TGFβ receptor II (dnTGFβRII)-armoured, prostate-specific membrane antigen (PSMA)-targeted CAR-T cells showed improved proliferation, persistence and antitumour efficacy in preclinical prostate cancer models (32). Early-phase testing subsequently demonstrated clinical feasibility and preliminary biological activity in metastatic castration-resistant prostate cancer, although grade ≥2 cytokine release syndrome with concurrent sepsis (33). Unlike dnTGFβRII, which confers intrinsic resistance to TGFβ signalling, secreted P144 can modulate the surrounding microenvironment. These approaches may therefore be complementary, potentially combining protection of the transferred cells with a relief of suppression in neighbouring immune populations. Other engineering strategies include base-editing dominant-negative mutations into the endogenous TGFβr2 locus and introducing switch receptors that convert TGFβ into a costimulatory signal (34–35). A complementary approach equips CAR-T cells to secrete microenvironment-modifying proteins. VEGF-neutralising scFv secretion improved activity across preclinical solid tumor models, while mesothelin-targeted CAR-T cells secreting an anti-PD-1 nanobody have entered first-in-human testing in mesothelioma (36–37).

Experience with protein therapeutics further illustrates the challenges of targeting this pathway. Bintrafusp alfa, a PD-L1 antibody fused to a TGFβRII trap, did not improve progression-free or overall survival over pembrolizumab in a randomised phase 3 trial in PD-L1-high advanced non-small cell lung cancer (38). More selective approaches include linavorkibart, which targets latent TGFβ1 and has shown a manageable safety profile with pembrozumab, and the GARP: TGFβ1-targeting antibody livmoniplimab (39–40). Collectively, these strategies seek either spatial restriction of TGFβ inhibition or selectivity for ligand activation and complex formation, reflecting the difficulty of inhibiting a pathway essential to tissue homeostasis and immune regulation (11,13). CAR-T-delivered P144TGFβ antagonist (in this case P144) offers a cell-based approach whose benefit may extend beyond direct protection of T cells to recruitment of innate effector cells. In vitro, P144 secretion increased cytokine production, particularly IFNγ, without measurably changing antigen-specific cytotoxicity. This dissociation is consistent with TGFβ-mediated suppression of effector cytokine production independently of target-cell lysis (41–42).

Local blockade may therefore primarily relieve a cytokine-production checkpoint rather than increase intrinsic cytolytic capacity. Because IFNγ regulates myeloid cell recruitment, activation and functional polarisation, Its increased production could contribute to contribute to the innate changes observed in vivo (43–44). Establishing this would require IFNγ neutralisation or genetic knockout.

The identification of neutrophils as functional contributors to P144-mediated efficacy is the principal mechanistic finding of the study. Using the same mesothelioma model, Frindlender and colleagues demonstrated that systemic TGFβ blockade shifted tumor-associated neutrophils (TANs) from a pro-tumorigenic N2 phenotype towards a hyper segmented, cytotoxic N1 phenotype, with neutrophils required for the antitumour effect (45). This work established the N1/N2 framework and identified TGFβ as a regulator of TAN plasticity (45–46). More recently, studies in lung carcinoma showed that Smad3 signalling maintains the N2 state: activation correlated with N2-skewed infiltrates and poorer survival in patients, whereas genetic depletion or pharmacological inhibition promoted antitumour phenotypes in experimental models (47). These findings support growing interest in exploiting TAN functional diversity therapeutically (48). Consistent with this framework, M11-P144 treatment increased intratumoral neutrophil accumulation and enhanced the ex-vivo cytotoxicity of tumor-derived neutrophils. However, we did not undertake the phenotypic or transcriptional analyses needed to establish equivalence to canonical N1 neutrophils and therefore distinguish the demonstrated recruitment and functional enhancement from the hypothesis that P144 induces N1-like polarisation.

The contribution of these cells to efficacy was tested directly. Anti-ly6G-mediated depletion substantially attenuated the advantage of m11-p144 over parental M11 CAR-T cells while having little effect on parental CAR-T activity. Neutrophils thus appear largely dispensable for baseline CAR-T-mediated control but become important when treatment includes P144 delivery, paralleling the earlier systemic TGFβ blockage findings (45). Depletion reduced rather than abolished the benefit, indicating that neutrophils account for only part of the response; increased car-t accumulation and cytokine production may provide additional contributions. This interpretation also accords with evidence that appropriately activated neutrophils can cooperate with T-cel-based immunotherapies and support tumor rejection (49–50).

Extension to FAP-targeted CAR-T cells demonstrates the modularity of the P144 secretion cassette. FAP-P144 T cells retained transduction efficiency, expansion and antigen-specific cytotoxicity while producing more IFNγ and achieving superior tumour control, accompanied by increased neutrophil accumulation within tumours. This is notable because FAP-directed cells primarily target tumor stroma rather than malignant cells. Stromal targeting can address extracellular matrix deposition, T cell exclusion and local immunosuppression, all major barriers to CAR-T cell efficacy in solid tumours (51). Previous studies by Wang et al and Lo et al showed that FAP-directed CAR-T cells can reduce desmoplasia, remodel the TME and enhance endogenous antitumour immunity (20,50). In pancreatic cancer models, this approach has also been proposed to facilitate subsequent tumor-antigen-directed therapy, including mesothelin-targeted CAR-T cells (19–20). Our findings suggest that P144 delivery can enhance both tumor and stroma-directed platforms, providing flexibility in addressing antigen-specific targets alongside stromal and cytokine-mediated resistance.

Clinically, mesothelin-targeted cellular therapies have established feasibility but shown limited efficacy. Regionally administered mesothelin car-t cells were safe and produced encouraging preliminary activity with anti-PD-1 blockade (53). Mesothelin-directed T cell receptor fusion constructs achieved a 20% objective response rate in Phase 1 testing, and anti-PD-1-nanobody-secreting mesothelin CAR-T cells produced responses in small first-in-human cohort (37, 54). Conversely, systemic mesothelin CAR-T cells produced no objective responses in advanced solid tumours, with stable disease as the best outcome and median progression-free survival of approximately two months (55). These results highlight barriers beyond antigen recognition, including restricted tumor access and suppression of effector function with desmoplastic, cytokine-rich lesions (19, 56).

Secretory peptide-based TGFβ blockade may address these barriers without modifying the CAR-T cells’ endogenous signalling machinery. All treatments in our study were well tolerated, with no weight loss or overt toxicity. Although these observations do not establish comparative safety, they support further evaluation of local delivery as an alternative to systemic pathway inhibition, whose development has been constrained by disruption of TGFβ-dependent homeostasis and immune regulation (11, 13, 57).

Several limitations warrant consideration in our study. First, it was designed and budgeted primarily to compare P144-secreting CAR-T cells with their parental counterparts; the pronounced neutrophil influx was unexpected. Within the time and finances, we had, we established its functional contribution but were unable to define the recruited cells’ exact phenotype, recruitment signals or cytotoxic mechanisms. Phenotypic and transcriptional profiling, chemokine analysis and further mechanistic studies are therefore near future priorities. The neutrophil depletion findings establish a contribution to efficacy independently of whether these cells ultimately meet canonical N1 criteria. Second, efficacy was evaluated in a single syngeneic mesothelioma model. Although AE17 reproduces important features of human mesothelioma, including a desmoplastic, TGFβ-rich TME, validation in additional models, including patient-derived xenografts, is needed. Third, differences between murine and human neutrophil biology leave uncertain whether human neutrophils will respond similarly to local TGFβ inhibition. Finally, the intratumoral concentration, diffusion radius and half-life of secreted P144 was not measured and will be important for further construct optimisation.

In summary, P144-secreting CAR-T cells improved mesothelioma control, accompanied by increased intratumoral CAR-T accumulation and cytokine output and recruitment of more cytotoxic neutrophils that contributed to the full therapeutic benefit. Activity across mesothelin- and FAP-targeted constructs supports platform modularity, although the recruited neutrophil population remains incompletely characterised. These findings support further preclinical and translational development of car-t-delivered TGFβ antagonists, for desmoplastic, immunosuppressive solid tumours.

## Data Availability Statement

The original contributions presented in the study are included in the article and Supplementary Material. Further enquiries can be directed to the corresponding author.

## Ethics Statement

All animal studies were approved by the institutional Animal Welfare Ethical Review Body (AWERB) and carried out under Project License PP9736772, in accordance with UK Home Office Regulations under the Animals (Scientific Procedures) Act 1986 and national and ARRIVE guidelines. Animals were housed in a pathogen-free facility in ventilated cages, at 20–24 °C and 40–60% relative humidity, with a 12 h light/dark cycle.

## Author Contributions

RB: Conceptualization, investigation, formal analysis, data curation, writing – review & editing; EH: Investigation, formal analysis. GB: Investigation. FS: Investigation, writing – review & editing. CLB: Investigation, writing – review & editing. RST: Investigation, writing – review & editing. VD: Investigation, writing – review & editing. RRDS: Conceptualization, investigation, formal analysis, writing – review & editing. AK: Conceptualization, methodology, supervision, funding acquisition, formal analysis, data curation, writing – original draft, writing – review & editing. All authors contributed to the article and approved the submitted version.

## Funding

This work has been funded through a Cris Cancer Foundation Fellowship grant, as well as a studentship grant by a private donor.

## Acknowledgements

We would like to thank Prof. Steven Albelda and the University of Pennsylvania for the valuable cell lines and plasmids. Special thanks to the Core Flow Cytometry facility and the Biological Services Unit at ICR for their advice and support throughout the project. This manuscript has been language-edited using Copilot for Microsoft Office 365.

## Conflict of Interest

The authors declare no conflict of interest.

## Supplementary Material

Supplementary Figure S1 is provided as a separate file.

## Supplementary Figure Legend

**Supplementary Figure S1. Baseline TGFβ production, tolerability, and broader immune profiling. (A)** TGFβ released into culture supernatant by HT1080, EMMESO, REN, AE17 and AB1 cells, measured by ELISA. **(B)** Body weight (g) over time for the study animals shown in Figure 2. **(C)** Frequency of CD45⁺, CD3⁺, CD8⁺ and CD4⁺ cells as a percentage of total live cells in the tumour digest across untreated (UT), non-transduced (NTD), M11 and M11-P144 groups. **(D)** Frequency of live CD45⁺, CD11b⁺, Ly6G⁺, Ly6C⁺, M1-polarised, M2-polarised and NK cells as a percentage of total live cells in the tumour digest across the same groups. All panels: mean ± SEM.

## Notes

### Competing Interest Statement

The authors have declared no competing interest.

