## Supplementary figures and images for "Anti-TGFβ peptide P144 augments CAR-T cell efficacy through neutrophil recruitment and TME remodeling in malignant mesothelioma"

### Supplementary Figure 1

Supplementary Figure 1.

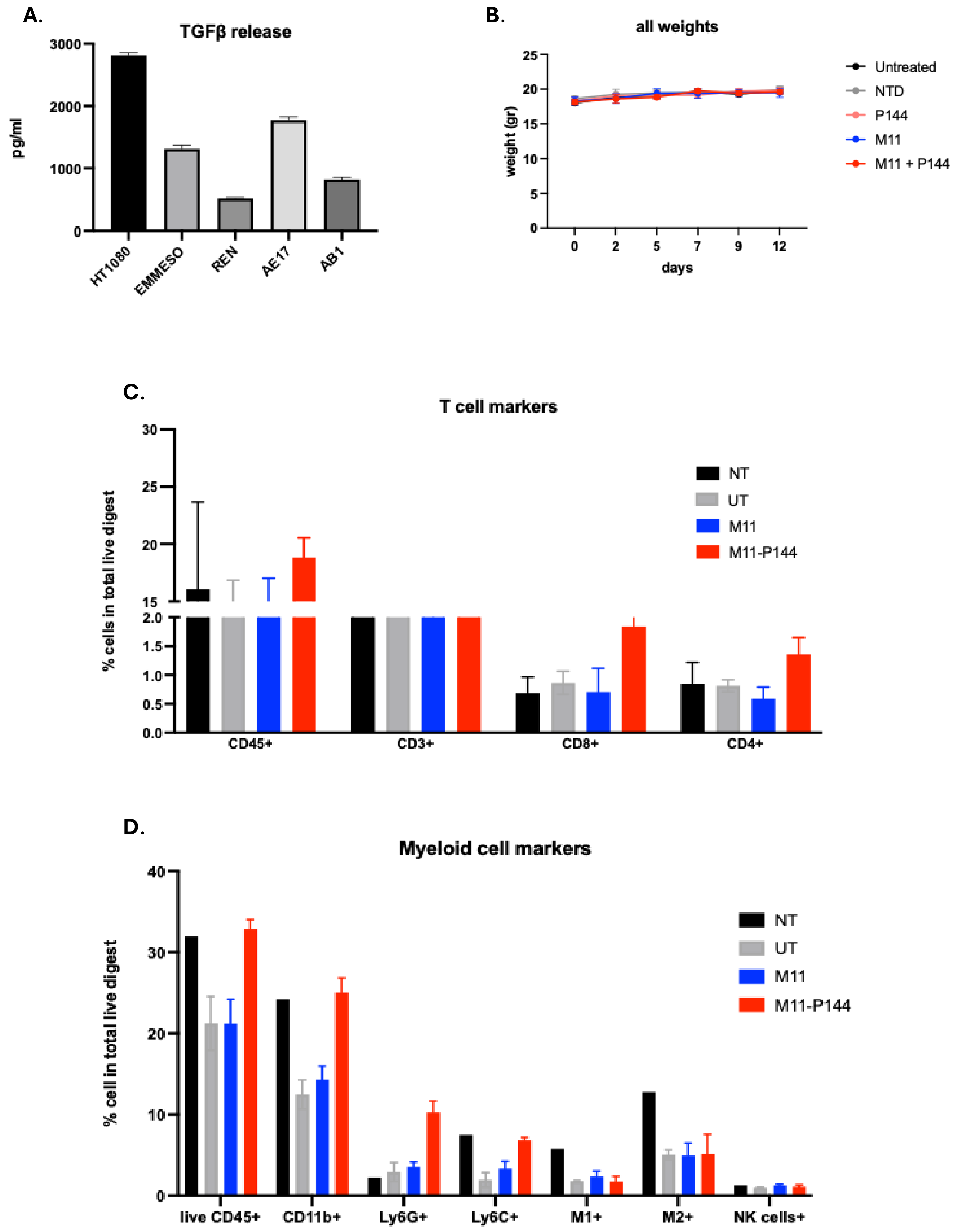
